# Allele segregation analysis of F_1_ hybrids between independent *Brassica* allohexaploid lineages

**DOI:** 10.1101/2021.11.18.469170

**Authors:** Daniela Quezada-Martinez, Jun Zou, Wenshan Zhang, Jinling Meng, Jacqueline Batley, Annaliese S. Mason

**Affiliations:** Plant Breeding Department, University of Bonn, 53115 Bonn, Germany; Plant Breeding Department, Justus Liebig University, 35392 Giessen, Germany; National Key Laboratory of Crop Genetic Improvement, Huazhong Agricultural University, Wuhan, China; School of Biological Sciences, The University of Western Australia, Crawley 6009, Australia

**Keywords:** *Brassica*, hexaploids, interspecific hybridization, karyotype rearrangement, homoeologous recombination

## Abstract

In the *Brassica* genus we find both diploid species (one genome) and allotetraploid species (two different genomes) but no naturally occurring hexaploid species (three different genomes, AABBCC). Although hexaploids can be produced via human intervention, these neo- polyploids have quite unstable genomes and usually suffer from severe genome reshuffling. Whether these genome rearrangements continue in later generations and whether genomic arrangements follow similar, reproducible patterns between different lines is still unknown. We crossed *Brassica* hexaploids resulting from different species combinations to produce five F_1_ hybrids, and analyzed the karyotypes of the parents and the F_1_ hybrids, as well as allele segregation in a resulting test-cross population via molecular karyotyping using SNP array genotyping. Although some genomic regions were found to be more likely to be duplicated, deleted or rearranged, a consensus pattern was not shared between genotypes. *Brassica* hexaploids had a high tolerance for fixed structural rearrangements, but which rearrangements occur and become fixed over many generations does not seem to show either strong reproducibility or to indicate selection for stability. On average, we observed 10 *de novo* chromosome rearrangements contributed almost equally from both parents to the F_1_ hybrids. At the same time, the F_1_ hybrid meiosis produced on average 8.6 new rearrangements. Hence, the increased heterozygosity in the F_1_ hybrid did not significantly improve genome stability in our hexaploid hybrids, and might have had the opposite effect. However, hybridization between lineages was readily achieved and may be exploited for future genetics and breeding purposes.

## INTRODUCTION

The *Brassica* genus belongs to the Brassicaceae family (Warwick et al., 2006) and includes many economically valuable crop species. The *Brassica* crops can be broadly classified into vegetable, oilseed, fodder and condiment types. During the 1930s, the chromosome number and genetic relationships between the cultivated *Brassica* species was established in what we know as the triangle of U (U, 1935): the diploid species *B. rapa* (AA, n = 10), *B. nigra* (BB, n = 8) and *B. oleracea* (CC, n = 9) were defined as progenitors of the allotetraploid species *B. juncea* (AABB, *n* = 18), *B. napus* (AACC, *n* = 19), and *B. carinata* (BBCC, *n* = 17), which originated via pairwise spontaneous hybridization between these diploids. The *Brassica* vegetables, *B. oleracea* and *B. rapa,* are characterized by vast diversity in subspecies and varieties (Cheng et al., 2016). In these species, it is possible to find different domesticated morphotypes distinguishable by leaf types, inflorescence types or enlargement of roots or stems. In *Brassica* species it is also possible to find traits of agronomic interest, such as disease resistance (Chevre et al., 1996; Fredua-Agyeman et al., 2019; J. Mei et al., 2011; Jiaqin Mei et al., 2013; Peng et al., 2014; Taylor et al., 2015), abiotic stress tolerance (Gill et al., 2011; Hayat et al., 2011; Irfan et al., 2014; Wilson et al., 2014) and pod shatter resistance (Y. Zhang et al., 2016), among other potential traits (reviewed in (Katche et al., 2019)).

Commonly, in the *Brassica* genus we find both diploid and tetraploid species but no naturally occurring hexaploid species (AABBCC = 2n = 6x = 54). Despite this, it is possible to synthesize this hybrid via human intervention. The three most common cross combinations to produce an allohexaploid are: (i) *B. carinata* × *B. rapa* (Figure 1A), (ii) *B. juncea* × *B. oleracea,* and (ii) *B. napus* × *B. nigra* (reviewed in (Gaebelein & Mason, 2018)), that from now on will be referred to as “carirapa”, “junleracea”, and “naponigra” allohexaploid types (Zou et al., 2010). All of these hybrids are usually colchicine-treated to induce chromosome doubling following hybridization between a diploid and a tetraploid species. A more recent method to produce an allohexaploid is via two-step crossing and relies on unreduced gamete production: the hybridization between *B. napus* × *B. carinata* × *B. juncea* (Mason et al., 2012), referred to as “NCJ” allohexaploid types (Figure 1A). Until now, many of the allohexaploid lines produced were solely used to cross to *B. napus,* either to introgress genetic diversity (Hu et al., 2019; Jiang et al., 2007; Li et al., 2004, 2006; Zou et al., 2010, 2018), or to transfer specific traits such as yellow seededness (Meng et al., 1998; Rahman, 2001) or fungal disease resistance (Sjödin & Glimelius, 1989). Despite this, a new allohexaploid hybrid has the potential to become a new species with the advantage of combining all the different traits present in the six Ús triangle species, thus broadening the genetic resources available for breeders (Chen et al., 2011). In such an allohexaploid, it would also be possible to take advantage of “fixed heterosis” (Abel et al., 2005), where the heterosis present between the subgenomes (A, B and C) can be maintained in inbreeding lines. In addition, new phenotypes which are not present in the original parents may develop from the different crosses via novel mutations due to the hybridization event (Kaur et al., 2014; Udall & Wendel, 2006).

**Figure 1.**
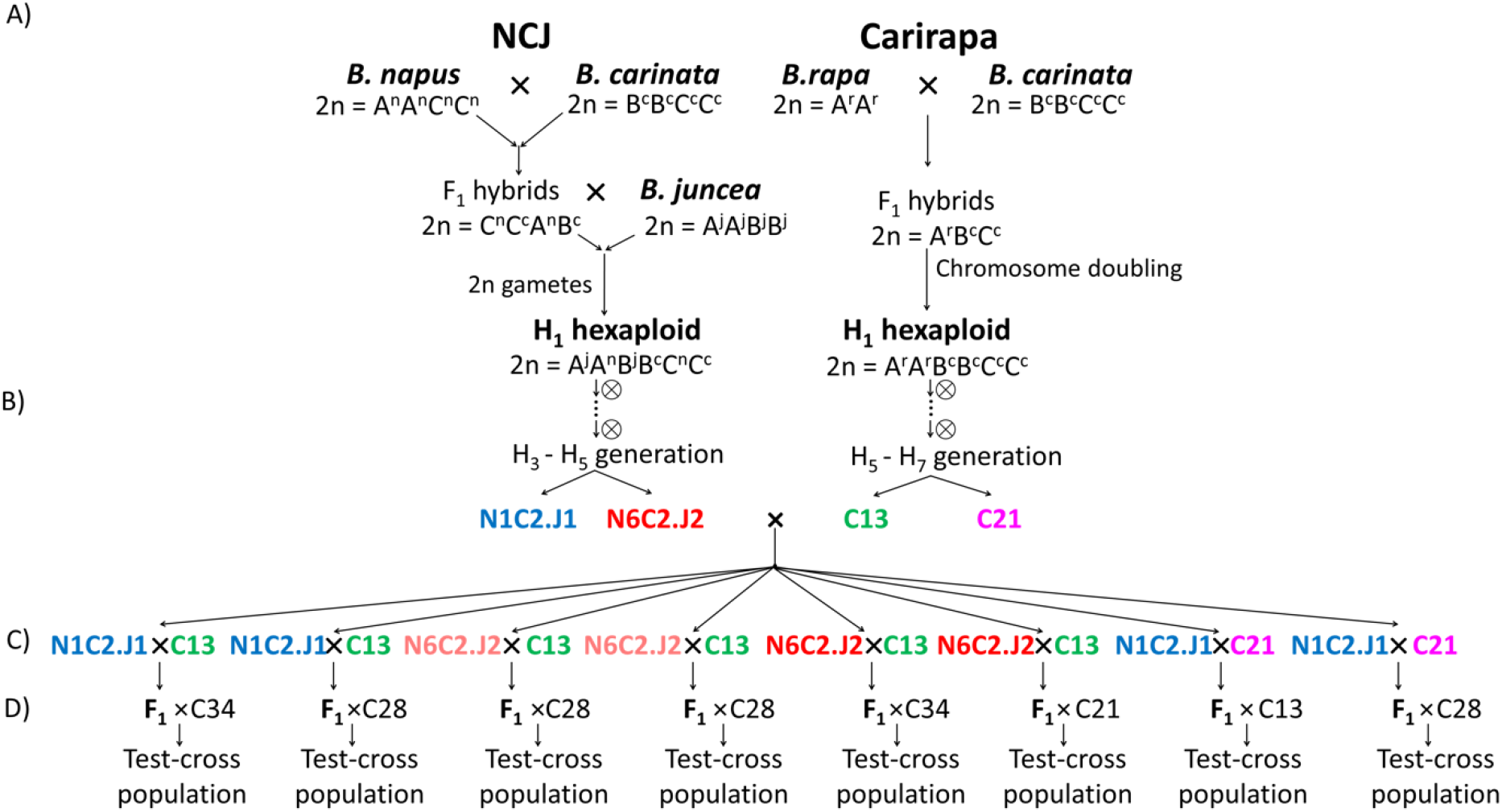
Crossing scheme. The arrows indicate the direction of the crossing. A) Crossings involved in the production of the NCJ and carirapa hexaploid lines. B) Selfing and selection of the genotypes. C) Crossings between NCJ and carirapa genotypes. For the NCJ genotype combination N6C2.J2 two different lineages were used for crossing, indicated by a different shade of red. D) F_1_ test-cross to another carirapa hexaploid.

Allohexaploid *Brassica* (AABBCC = 2n = 54), as a new polyploid has to overcome the major challenge of establishing regular meiosis (Pelé et al., 2018). The correct pairing of homologous chromosomes during meiosis is a key factor to ensure correct cross-over and chromosome segregation. In the case of this allohexaploid, meiosis is challenging due to the presence of three ancestrally homologous chromosome sets from different evolutionary lineages, known as homoeologous chromosomes (Ramsey and Schemske 2002). If non-homologous pairing during meiosis occurs, it can lead to different chromosomal rearrangements, such as deletions, duplications and translocations (Udall et al., 2005), heavily affecting genome stability. In a synthetic trigenomic hybrid, we usually observe the A and the C genome are more likely to pair during meiosis, compared to the A-B or C-B homoeologs (Mason et al., 2010). In the case of natural allopolyploids, such a *B. napus,* meiosis progresses normally, although non-homologous exchanges do still occur at lower frequency (Higgins et al., 2018; Parkin et al., 1995; Udall et al., 2005).

Many different types of cross-combinations have been tested to produce meiotically stable 2n = AABBCC allohexaploids (reviewed in (Gaebelein & Mason, 2018)). It has been shown that early generation (F_2_) carirapa hexaploids have low pollen viability and irregular configurations during meiosis (Howard, 1942; Iwasa, 1964). It has also been seen that fertility and number of 2n = 54 (putatively euploid) plants can increase by selection over successive generations (Tian et al., 2010; Zhou et al., 2016). Genotype combinations seem to play a great role in the success of the hexaploid, as many of the more stable and fertile hexaploids produced are derived from just a few lines (Tian et al., 2010). A specific *B. rapa* genotype (R01) was also reported to result in a meiotically stable carirapa allohexaploid (Gupta et al., 2016). Carirapa combinations that included this *B. rapa* genotype had high frequency of bivalent formation during meiosis, progeny with the expected 54 chromosomes, and no major translocations observed (Gupta et al., 2016). Similarly, genotype-specific effects on fertility and meiotic stability have been observed in NCJ hexaploids (Mwathi et al., 2017).

In the present study we analyzed chromosome and allele segregation resulting from the meiosis of five different F_1_ hybrids produced between crosses of putatively stable *Brassica* hexaploids carirapa and NCJ. We hypothesized that the new hybrids might show improved meiotic stability (more regular chromosome segregation and fewer non-homologous recombination events) compared to their parents. We also aimed to determine if chromosome rearrangements found at high frequency or in independent lineages of the parent hexaploid types (e.g. both in NCJ and carirapa) might be associated with improved meiotic stability, since all lineages underwent strong selective pressure for fertility.

## MATERIALS AND METHODS

### Plant material and crossing scheme

The most advanced and fertile lines available were from the two NCJ parental genotype combinations N1C2.J1 and N6C2.J2 (Mason et al., 2012; Mwathi et al., 2017) and hence one and two lineages from these genotype combinations, as well as one lineage from carirapa allohexaploid genotypes (C13, C21) (Tian et al., 2010), were selected based on chromosome number and total seeds produced (Figure 1B, Supplementary Table 1). Initially, two plants from the hexaploid genotypes C13 and N6C2.J2 were grown and crossed under greenhouse conditions at Justus Liebig University, Giessen, Germany, while the rest of the lines were grown under field conditions at Huazhong Agricultural University, Wuhan, China. In the first round of crossings (F_1_ hybrid production), the NCJ hexaploid type was used as a female parent, and the carirapa parent was used as the pollen donor (Figure 1C). The cross was done by hand, via emasculating flower buds and gently rubbing the anthers over the exposed stigma. The pollinated buds were then labeled and covered with a microperforated bag to prevent contamination from other pollen sources. The F_1_ hybrid seeds were collected and grown under field conditions at Huazhong Agricultural University, Wuhan, China. To be able to analyze the meiotic performance and allele segregation from this new F_1_ hybrid, a test-cross was carried out. A total of eight F_1_ hybrid plants were selected and test-crossed to another carirapa hexaploid (genotypes C21, C28 and C34) (Tian et al., 2010) (Figure 1D). The test cross-progeny were then grown under field conditions at Huazhong Agricultural University, Wuhan, China.

### DNA extraction and SNP analysis

From each plant, a piece of leaf sample was collected and DNA was extracted using the CTAB method (Doyle & Doyle, 1990). For some of the plants, leaf sample from sibling lines was used as no leaf sample was stored from the original line. The DNA was then genotyped using the Illumina Infinium *Brassica* 90K SNP array (Ilumina, San Diego, CA, USA) following manufactureŕs instructions. A total of 77 970 SNPs distributed across the A (23 482), B (25 822), C (26 731), and unplaced location (1 877) *Brassica* subgenomes were obtained after applying the recommended cluster file for the A and C genomes (Clarke et al. 2016, Supplementary Data set Table 2) and by automated clustering in Genome Studio for the B genome. SNP positions for the A and C genomes were determined by the top hit (highest *e*- value) based on BLAST to the *B. napus* Darmor-*bzh* v. 8.1 reference genome (Bayer et al., 2017). For the B genome, we used the positions provided in the Illumina Infinium 90K SNP array, which were based on the *B. nigra* Ni100 short read reference genome assembly (Perumal et al., 2020). The SNP data was initially cleaned by removing non-specific alleles and SNPs with undetermined genomic locations.

The first step in the analysis was to carry out paternity testing to verify the genetic composition of the F_1_ hybrid between the NCJ and carirapa allohexaploid parents for each of the eight populations. For this, homozygous polymorphic alleles for each parent in the B subgenome were used as diagnostics. If the expected heterozygosity was observed in the F_1_ line (e.g. NCJ allele AA × carirapa allele BB → F_1_ hybrid allele AB) it was considered a true F_1_ hybrid. The second step was to determine true segregating progeny between the F_1_ hybrid and the test- cross carirapa allohexaploid parent. To do this, the same approach was used by selecting homozygous polymorphic alleles in the B subgenome between the F_1_ and test-cross carirapa parent: if the segregating progeny had the expected heterozygous allele, they were considered true test-cross progeny. If the population were true hybrids in both analyzed cases, subsequent analyses were done.

### Chromosome count and molecular karyotyping

To determine chromosome presence or absence we used SNP and Log_2_ R ratio data (Supplementary Data set Table 3). The absence of both chromosomes was seen across the entire chromosome, or most of it, as “no call” SNPs (NC). If the chromosome was present in the SNP data, the Log_2_ R ratio data was used to determine loss or gain of a chromosome copy by assessing the inheritance of the centromeric region of each chromosome (inheritance of the centromere was inferred to mean inheritance of that chromosome, since chromosome fragments cannot be transmitted without a centromere). Experimentally-derived values were used based on comparison to hybrid standards with known haploid and diploid chromosome complements. Log_2_ R ratios between -0.5 and -0.2 were assumed to indicate one missing chromosome copy, Log_2_ R ratios between 0.2 to 0.5 were assumed to indicate gain of an extra copy and Log_2_ R ratios of more than 0.5 to indicate more than one extra chromosome copy gained. The approximate centromere locations for the A and C chromosomes were established according to the *B. napus* Darmor-*bzh* v8.1 reference genome (Bayer et al., 2017), based on remapping of previous genetic data (Mason et al. 2016). For the molecular karyotyping, a similar approach was used. If no-call (NC) SNPs covering ≥ 1 mega bases (Mb) in the chromosome was observed, it was categorized as both copies missing. A similar size cut-off was used for the Log_2_ R ratio values. Missing regions covering less than 1 Mb were not considered in the analysis due to consideration of SNP distribution and density constraints. The translocations between the genomes were established based on homoeology relationships between the A, B and C genomes (Chalhoub et al., 2014; Perumal et al., 2020). The final karyotype was plotted in RStudio using the R package chromDraw (Janečka & Lysak, 2016) and posterior editing was done in GIMP ver. 2.10.20.

### Allele segregation in the F_1_ hybrid

To analyze the allele segregation from the F_1_ hybrid, the alleles from each parent were analyzed independently. To analyze the NCJ parent, homozygous polymorphic alleles from the NCJ vs. carirapa parent 1/carirapa test-cross parent were filtered (NCJ allele: AA vs. carirapa parent 1/carirapa test-cross parent: BB), and vice versa for the carirapa parent. In each of the eight populations, the number of test-cross progeny ranged from 14 – 22 (Table 1). The 1 : 1 AB : BB observed vs. expected allele segregation was tested using a Х^2^ test with a significance level of p < 0.05.

**Table 1.**
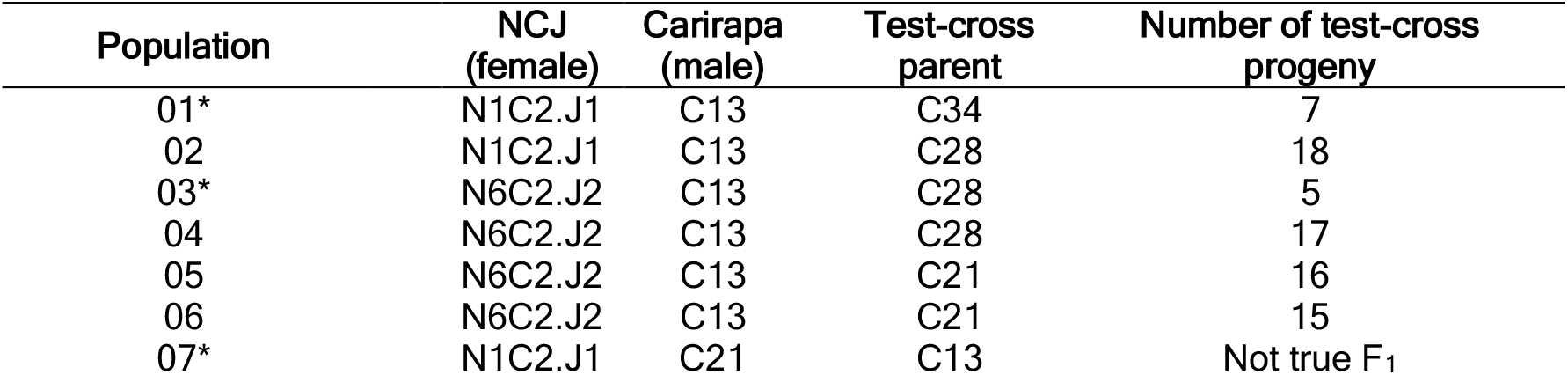
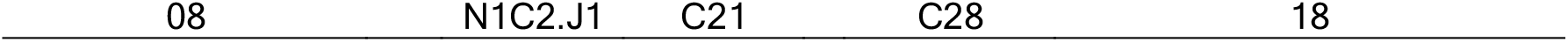
Brassica allohexaploid populations. Genotypes crossed between carirapa (C13 and C21, S5- S7) and NCJ (N1C2.J1 and N6C2.J2, S3-S5) allohexaploid types. Test-cross progeny correspond to the total number of progeny that were determined to comprise true test-cross progeny between the F_1_ and the test-cross parent. *Populations removed from further analyses.

### Data Availability Statement

Supplemental data files are available at FigShare. File S1 contains a complete description of the plant material, File S2 contains the genotyping data (allele calls) from the Illumina Infinium *Brassica* 90K SNP array, and File S3 contains the Log R Ratios for each SNP based on the Illumina Infinium *Brassica* 90K SNP array.

## RESULTS

### Pre-existing fixed translocations in *Brassica* parental genotypes

*Brassica* genotypes used to produce the carirapa and NCJ allohexaploids were analyzed for fixed rearrangements in the genome. The parental lines used to produce the carirapa hexaploids, *B. carinata* accession “03949”, and the *B. rapa* genotypes “Ankangzhong” and “WulitianYC” did not have any detectable fixed translocation events in their corresponding genomes. In the case of the lines used in the crossing of NCJ hexaploids, we only observed fixed events in the *B. napus* genotypes. In the case of the *B. napus* genotype “Surpass400_024DH” (N1) we detected deletions in chromosomes A01 and A04, located at 2.2-4.7 Mb and 20.7-23-3Mb, respectively. In the C genome of “Surpass400_024DH”, we identified a deletion at the top of chromosome C01, located between 1.4-3.3 Mb. We also found two other deletion events located in chromosome C09, at positions 0-2 and 52.9-60.1 Mb, respectively. For the “Ag-Spectrum” genotype (N6) we observed a deletion on chromosome C02, located between 8-10.7 Mb.

### Chromosome number in NCJ and carirapa *Brassica* hexaploids

From the available NCJ and carirapa collection, the most advanced and fertile lines were selected. From the NCJ type, the genotypes selected were N1C2.J1 and N6C2.J2 and from the carirapa type, C13 and C21. The carirapa lines were not fully homozygous, suggesting that pollen cross-contamination with other genotypes may have occurred during propagation in the field. In the NCJ-type hybrids, residual heterozygosity was present based on the method of producing these hybrids, but no cross-contamination was indicated by the presence of alleles other than those from the parent genotypes. In total, six plants per allohexaploid type were grown and later on crossed to produce F_1_ hybrids between the lines.

First, the relative chromosome number for the 12 allohexaploid parental lines was determined using SNP data and Log_2_ R ratio values (Figure 2). For the “carirapa” type, the chromosome numbers were 51 (1 plant), 52 (2 plants), 53 (1 plant) and 54 (2 plants). In the case of the allohexaploid NCJ, the chromosome numbers observed were 50 (1 plant), 51 (1 plant), 52 (1 plant), 53 (1 plant), 54 (1 plant) and 55 (1 plant). The two carirapa lines with 54 chromosomes were aneuploid in terms of genome composition. In contrast, the NCJ line with 54 chromosomes was a euploid line (genotype combination N1C2.J1). Overall, in both hexaploid types, the chromosomes A05, A06, A08, A10, C03, C05, C07, B01, B02, B04, B06 and B08 showed no changes (Figure 2). Specifically, in the carirapa lines, no B chromosome changes were observed. Changes of B chromosomes were however observed in the NCJ lines, for chromosomes B03 (two lines), B05 (two lines) and B07 (one line). For both lines, most of the chromosome number changes were observed in A01-A03, C01 and C02 chromosomes. Also, the chromosomes A01, B05 and B07 were lost in one carirapa line and in two independent NCJ lines.

**Figure 2.**
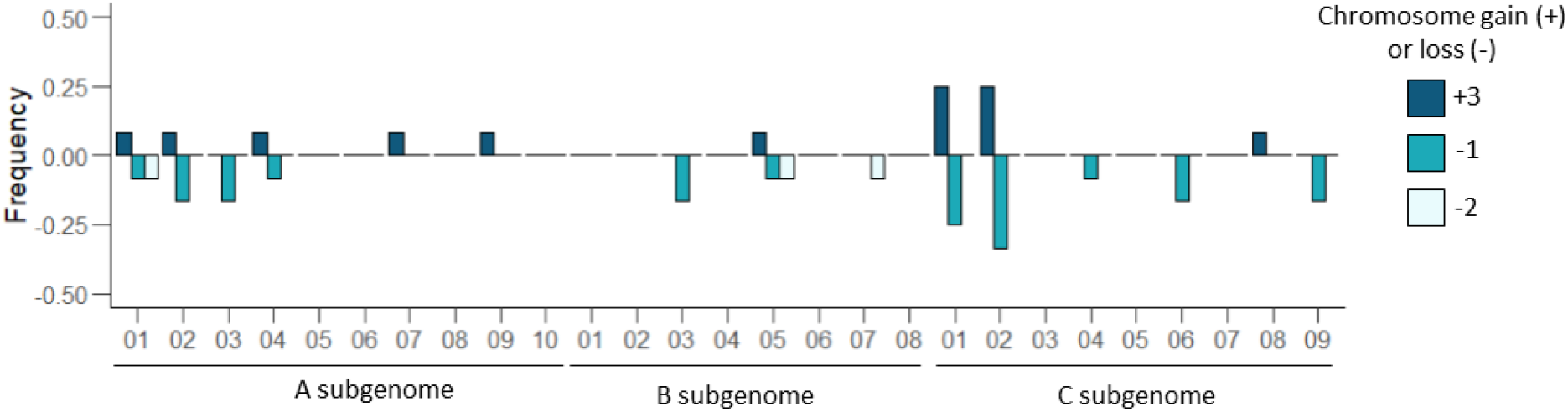
Frequency of chromosome variation in allohexaploid carirapa (*Brassica carinata* × *Brassica rapa*) and NCJ (*Brassica napus* × *Brassica carinata* × *Brassica juncea*) allohexaploid types. Whole chromosome changes (gain or loss) in 12 different allohexaploid plants across the three *Brassica* genomes (A, B and C).

### Fixed chromosome translocations in NCJ and carirapa allohexaploid types

As the NCJ and carirapa allohexaploids have been independently selected by fertility (total seed number) through several generations, we analyzed the presence of fixed translocations between the genomes (Figure 3). A single fixed event involving the B genome was detected between B01/A04, where the B01 segment was lost (deletion) and replaced by an A04 segment (duplication) for this homoeologous region. In other lines, the end of A04 (∼ 3Mb) was frequently lost, as was the case in seven out of 12 allohexaploid parents (four carirapa and three NCJ). The deletion event was already present in the *B. napus* cv. “Surpass400_024DH” used in the cross, and was inherited and fixed in the three NCJ plants analyzed.

**Figure 3.**
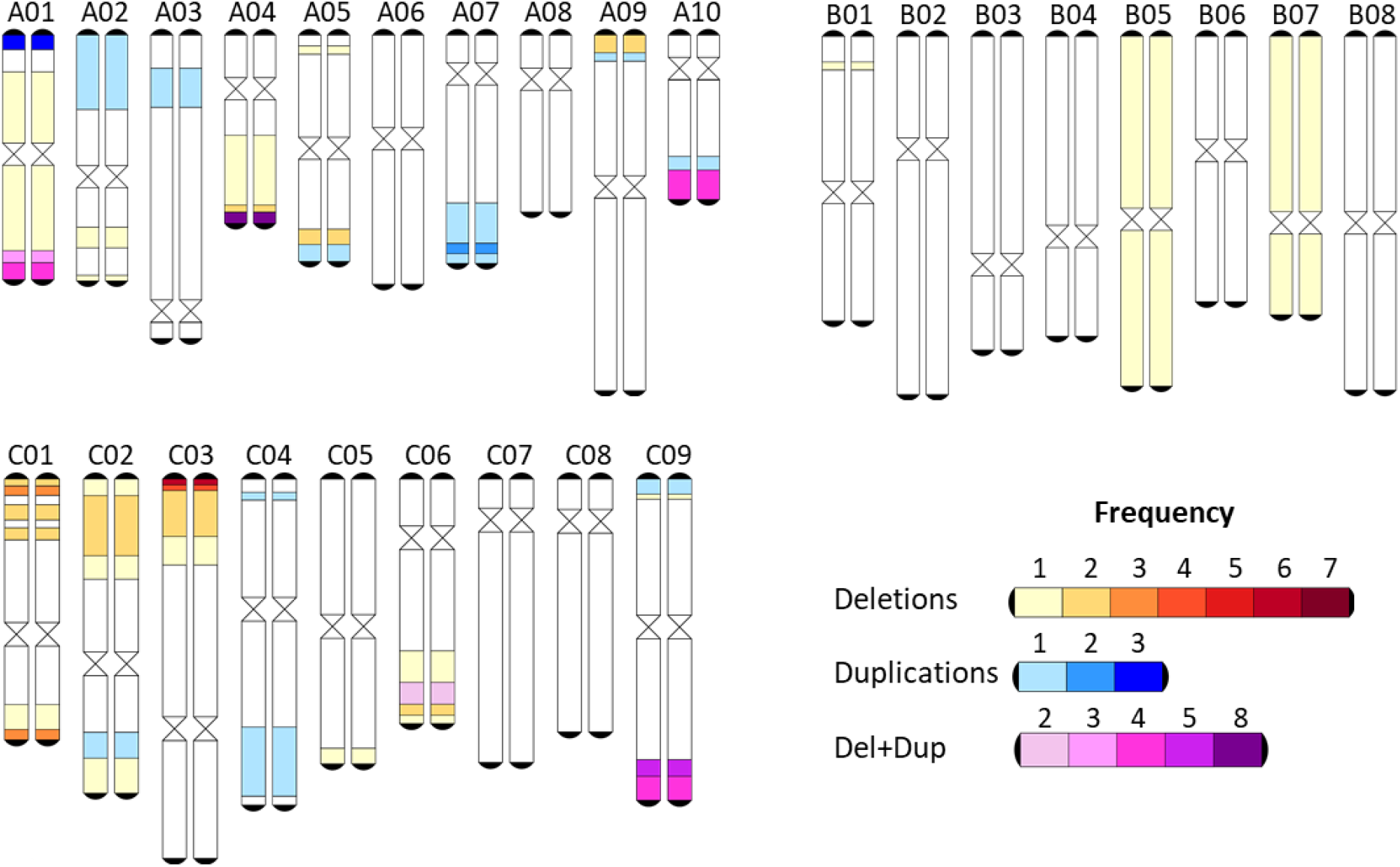
Karyotype of fixed events in *Brassica* allohexaploids NCJ (*B. napus* × *B. carinata* × *B. juncea*) and carirapa (*B. carinata* × *B. rapa*) type. The fixed partial or whole chromosome events are colored according to the legend for each of the three *Brassica* genomes.

For chromosomes B05 and B07, both copies were missing in two independent N6C2.J2 lineages, respectively. In the case of the carirapa type, the region was most likely replaced by the end of chromosome C04, but evidence of this was inconclusive in the NCJ types. Another region that was frequently lost was located at the top of chromosome C03, ranging from approximately 1.9 – 2.3 Mb in size: this region was missing in all four carirapa C13 parents (representing two independent lineages). The three plants from the genotype combination N1C2.J1 (one lineage) had in common a duplication/deletion event involving an extra copy of the top of chromosome A01 (∼ 2 Mb) being translocated into chromosome C01. The deletion of C01 was already present in the *Brassica napus* cv. “Surpass400_024DH” (N1) used to produce the hexaploid, unlike the duplication of A01 which was a new translocation event. In some regions, we found the overlap of events (duplications and deletions). The region at the end of chromosome A01 was duplicated in one lineage of the carirapa C13 (two parents), deleted in one carirapa parent (C21), and deleted in one NCJ parent (N6C2.J2). In the homoeologous region, there was no detectable duplication for any of the lines, hence it was more likely to be only a deletion event. Another region of overlap was found for homoeologous translocations between chromosomes A10 and C09. The end of A10 had three duplication events and one deletion event, while C09 had one duplication and four deletions located at ∼ 51-55 Mb; for some lines this deletion extended to the end of the chromosome (∼ 55-60 Mb). For the NCJ plants from the genotype combination N1C2.J1, the translocation between C09/A10 was already present in the *B. napus* cv. “Surpass400_024DH”, and was inherited and fixed in the hexaploids. The rest of the events were more line-independent (occurred in only one lineage). In total, we observed 50 fixed deletion events (32 carirapa, 18 NCJ) and 33 fixed duplication events (23 carirapa and 10 NCJ) involving the A, B and C genome of *Brassica* hexaploids. The fixed events observed in the carirapa hexaploids were not identified in either the *B. carinata* or the *B. rapa* accessions used as parents.

### Crossing NCJ and carirapa allohexaploids: F_1_ hybrid and allele segregation

Seven out of eight F_1_ hybrids were the result of a cross between carirapa and NCJ hexaploids, as expected (Table 1); one hybrid appeared to result from an outcross to an unknown paternal plant. The second step was to analyze the cross between the F_1_ hybrid and the test-cross parent. To do this, the same approach as in the F_1_ hybrid was used. In this case, populations 01 and 03 had only 7 and 5 individuals that corresponded to true test-cross progeny and the remaining individuals resulted from unintended self-pollination to produce F_2_ progeny seeds. The other five populations were used to analyze allele segregation from the F_1_ hybrid.

### Allele segregation per population

#### Population 02

The F_1_ hybrid was the result of a combination between the genotypes N1C2.J1 × C13. This hybrid had 52 chromosomes distributed between the A genome (19), B genome (16) and C genome (17) (Figure 4A). In the A genome, the chromosomes A01 and A05 were present in only a single copy, with just the chromosome from the carirapa parent present. Chromosome A01 was already a single chromosome in the NCJ parent, unlike A05, where both copies were present in the parent. The other chromosomes (A02-A09) were present in two copies, while chromosome A10 had at least one extra copy from the carirapa parent. In the A genome, non-reciprocal translocations between A05/C04, A05/C05, A09/C09 and A10/C09 were present in the chromosomes inherited from the carirapa parent. No translocations were observed in A-genome chromosomes from the NCJ parent. The bottom of chromosome A04 from NCJ was missing 2.5 Mb, corresponding to a known deletion present in the *B. napus* cv. “Surpass400_024DH” used as a parent in the crossing. The top of chromosome A09 from the NCJ parent was missing ∼ 3Mb that corresponded partially to an extended deletion that was initially inherited from a deletion also present in the *B. napus* cv. “Surpass400_024DH”. For the B genome, the F_1_ hybrid contained the expected 16 chromosomes with no translocations or deleted regions detected. In the C genome, the chromosomes C01-C08 were correctly inherited from the carirapa parent. In the case of chromosome C09, just a portion (∼5 Mb) from the end of the chromosome was present, as a translocated region into chromosome A10. Part of the top of chromosome C03 (∼7 Mb) was lost from the carirapa parent. The chromosomes C01-C09 were correctly inherited from the NCJ parent, despite the presence of only one copy of chromosome C02 in the parent. Translocations were observed in chromosomes coming from both parents. In the case of the carirapa parent chromosomes, the translocations present in the F_1_ were C01/A01 and C02/A02, and in the NCJ parent chromosomes there were C01/A01, C05/A05, C06/A07, and C09/A10. From the above translocations, only six corresponded to *de novo* events identified in the F_1_ hybrid.

**Figure 4.**
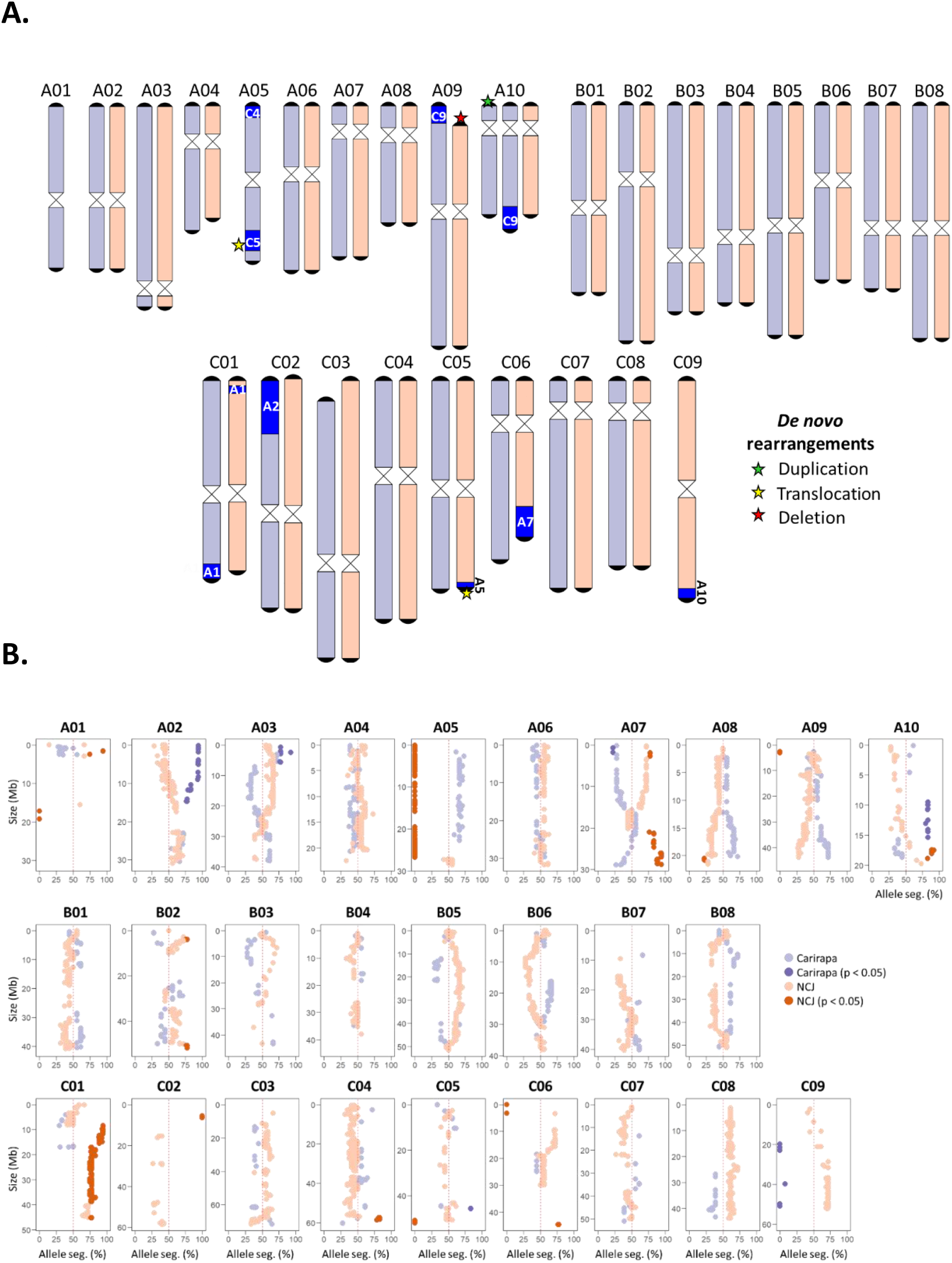
A. Molecular karyotype and B. allele segregation for F_1_ hybrid population 02. Chromosomes are colored based on hexaploid parent: carirapa in purple, NCJ in orange. Rearrangements are colored in blue. *De novo* translocations (present in the F_1_ hybrid but not in the parents) are marked with a star in a different color depending on the type (see legend). Chromosome sizes are represented in megabases (Mb). Expected segregation ratio of the alleles (50%) is marked with a red dotted line for each chromosome. Significant allele distortion (Х^2^ test, p < 0.05) is indicated with dark orange (NCJ) or dark purple (carirapa). Seg. = segregation.

Allele inheritance from the F_1_ hybrid into the test-cross progeny was also analyzed (Figure 4B). Segregation distortion was established if the observed allele segregation ratio was significantly different from the expected (Х^2^ test, p < 0.05). This test was performed independently for each of the parents of the F_1_ hybrid. In population 02, 18 test-cross progeny individuals were used to assess allele segregation (for more details see materials and methods). The allele segregation from the A01 and C02 chromosomes could not be established due to the lack of enough polymorphic alleles. The segregation distortion observed in the chromosomes A05 and C09 corresponded directly to the presence of only a single chromosome in the carirapa or NCJ parent, respectively. The allele segregation distortion observed in A07, A09, A10, and C05 corresponded to translocations present in the F_1_ hybrid. Allele segregation distortion not directly explained by the karyotype of the F_1_ parent was observed in chromosomes A02 (extension of a known duplication present in the F_1_ hybrid, carirapa), A03 (although few markers were represented, carirapa), C01 (NCJ), C04 (NCJ) and B02 (NCJ). Potentially, these events correspond to *de novo* rearrangements produced during meiosis in the F_1_.

#### Population 04

The F_1_ hybrid resulting from the combination between NCJ N6C2.J2 × carirapa C13 had 51 chromosomes distributed between the A (21), B (15) and C (15) genomes (Supplementary Figure 1A). In the F_1_ hybrid, chromosomes A02 and A07 had at least one extra copy from the carirapa and from the NCJ parent, respectively. These chromosomes were also doubled in the corresponding allohexaploid parent. Chromosome A04 was present as a single copy in the F_1_ hybrid, inherited only from the NCJ parent. In the carirapa parent, chromosome A04 was present as a single copy and it was not inherited to the F_1_. In the F_1_, the chromosome A05 inherited from the carirapa parent had a small region missing (1.6 Mb) located at chromosome position ∼24 Mb, and no potential candidate translocation corresponding to this region was identified. In the B genome, the F_1_ hybrid had only one B07 chromosome (carirapa origin). Chromosome B07 was completely absent in the NCJ parent. In the B03 chromosome of the F_1_ hybrid (NCJ origin) a duplicated region was identified, but it was not possible to determine the position in the genome of this extra copy (colored gray, see Supplementary Figure 1A).

In the C genome, chromosomes C01, C02 and C06 were present in single copies (NCJ). The chromosomes C01 and C02 were also present as a single copy in the carirapa parent, unlike C06, which was present as two copies in the carirapa parent and as a single copy in the NCJ parent. In the A genome, translocations between homoeologs A01/C01, A02/C02, A07/C07 and A09/C08 were observed. In the B genome just one translocation between B01/A04 (NCJ) was observed. In the C genome, we had translocations involving C02/A02, C04/A04, C05/B01 and C07/B02.

In the analysis of the allele segregation, all significant distortions were explained by the translocations described above (Supplementary Figure 1B). The exceptions were present in chromosomes A02 (carirapa), A07 (carirapa), A10 (carirapa), C06 (NCJ), B01 (carirapa) and B02 (NCJ). In addition, the single B07 chromosome in the F_1_ hybrid was expected to be present in 50% of the test-cross population, but was only present in 18% of the individuals (3 out of 17).

#### Population 05

The F_1_ hybrid (N6C2.J2 × C13) had 54 chromosomes distributed between the A (21), B (15) and C (18) genomes (Supplementary Figure 2A). Chromosomes with at least one extra copy were present for A04 (NCJ) and C03 (NCJ). Chromosome A04 was already doubled in the NCJ parent, unlike chromosome C03, which resulted from a new chromosome duplication event. Single copies were observed for chromosomes B05 (carirapa) and C09 (NCJ). In the case of B05, both copies were missing in the NCJ parent. Chromosome C09 was present as a single copy in the carirapa parent, and did not get inherited into the F_1_ hybrid. In the chromosomes from NCJ, translocations between A01/C01, A03/C03, A04/C04, B01/A05, C03/A03, and C08/A09 were found. In the case of chromosomes coming from the carirapa parent, we observed translocations involving the chromosomes A02/C02, A05/C04, A09/C09, C01/A01, C02/A10 and C03/A03 (Supplementary Figure 2A).

In the population segregating for alleles from the F_1_ hybrid, most of the allelic distortion was explained by CNV events in the parent F_1_ (Supplementary Figure 2B). The exceptions to this (putatively novel CNV events) were located in A02 (carirapa), A08 (NCJ and carirapa), B03 (NCJ), B04 (NCJ), B07 (NCJ), C02 (NCJ), C04 (NCJ and carirapa) and C08 (NCJ). Chromosome C09, as mentioned before, had just one copy in the F_1_ hybrid and it was present in fewer test- cross individuals than expected (25% presence vs. 50% expected). In the case of B05 as a single chromosome, only a few polymorphic alleles were present with which to make a proper comparison, but it also seemed to be present less often than expected (25% presence vs. 50% expected).

#### Population 06

The F_1_ hybrid in this population was a cross between N6C2.J2 × C13 and had 51 chromosomes distributed between the A (19), B (15) and C (17) genomes (Supplementary Figure 3A). Single copy chromosomes were A04, B05 and C08. Chromosome B05 was already missing both copies in the NCJ parent. In the case of A04 and C08, both copies were present in the corresponding carirapa and NCJ parents, respectively. No extra chromosomes were observed. Translocations between A01/C01 and A09/C08 were detected in NCJ chromosomes. In the case of the carirapa chromosomes, observed translocations were located between A09/C09- C08, C01/A01 and C02/A02.

When analyzing the allele segregation for alleles from the F_1_ hybrid (Supplementary Figure 3B), most of the distortion was explained by rearrangement events, with the exception of the following: end of A02 (carirapa), A03 (carirapa and NCJ), end of chromosome A04 (NCJ), A09 (carirapa), C01 (NCJ), C04 (NCJ), and C09 (NCJ). The A09 chromosome was a special case, as it was present in two copies in the F_1_ hybrid, but in the test-cross population the A09 of carirapa origin was present at a higher frequency than expected (expected 50%, observed 86.6%). Interestingly, this chromosome had two translocations involving chromosomes C9 and C8 c. The single chromosomes A04 and C08 segregated as expected in the population, unlike B05, which was present in only two out of the 15 test-cross population individuals and not in half of them, as expected. No other changes were observed in the B genome.

#### Population 08

The cross between N1C2.J1 (euploid, 2n = 54) × C21 gave rise to an F_1_ hybrid with 51 chromosomes distributed between the A (19), B (16) and C (17) genomes (Supplementary Figure 4A). Single chromosomes were observed for A03 and C01. Both chromosomes were present as two copies in the corresponding NCJ and carirapa parents. No chromosomes were doubled. In the NCJ chromosomes, we observed translocations between A02/C02, C01/A01, C06/A07, and C09/A10. Chromosomes A04 and A09 inherited from the NCJ parent had a deletion at the bottom and at the top of the chromosome respectively. These deletions were already present in the parents and we did not observe a duplicated homoeologous region that could have replaced this fragment in the F_1_ hybrid. In the chromosomes from the carirapa parent, we observed translocations between A01/C01, C02/A02, C03/A03 and C09/A09.

In the allele segregation from the F_1_ hybrid (Supplementary Figure 4B), most of the events were explained by the rearrangements, with the exception of areas on chromosomes A03 (NCJ), A06 (carirapa), B02 (NCJ), B06 (NCJ), B08 (NCJ and carirapa), C08 (NCJ) and C09 (NCJ).

### *De novo* rearrangements and inheritance in the F_1_ hybrids

Overall, 50 new rearrangement events (22 duplications and 28 deletions) were observed in the F_1_ hybrids. Out of these events, 28% (6 duplications and 8 deletions) were triggered by a previous event nearby or overlapping the chromosomic location of the new event already present in the hexaploid parent. On average, there were 10 new events per population, affecting mostly the A (52%) and C genome (44%). Both parents contributed almost equally to the *de novo* rearrangements observed in the F_1_ hybrids, with the exception of population 5, where the NCJ parent contributed to 10 rearrangements compared to 4 coming from the carirapa parent. As mentioned before, the least affected genome was the B, where just two events in two populations were observed. In the A genome, the chromosome most affected by rearrangements was A09, with a total of 8 events (2 duplications and 6 deletions). In the case of the C genome, the chromosome with more *de novo* rearrangements was C02, with six events (2 duplications and 4 deletions). We also observed nine *de novo* events involving whole chromosomes, where six chromosomes were lost and three were present in an extra copy in the F_1_ hybrid.

In the F_1_ hybrids, we also observed that some of the rearrangement events present in the parental hexaploid lines were either inherited in the same size or reduced in size due to cross- overs. In total, 72 rearrangements were inherited from the hexaploid parental lines with an average of 14.4 rearrangement events per F_1_ hybrid. Out of the 72 rearrangements, 8.3 % had a reduction in size due to a cross-over. Most of the events inherited from the parents corresponded to deletions (66.7 %), with seven events involving the loss of a chromosome, affecting mostly the C genome (27 events).

When analyzing the putatively *de novo* events produced by the F_1_ hybrid, we observed a total of 43 events (8.6 events on average per population). Out of these events in the F_1_ hybrid, 25 events were potential duplications and 18 deletions. Interestingly, many of these events also affected the B genome (9 events total, 8 from NCJ and 1 from carirapa origin, respectively). Overall, more events were produced by one meiosis in the F_1_ hybrid than by meioses coming from the grandparents, although the difference was only significant between the carirapa parent and the F1s (Figure 5, one way-ANOVA, p < 0.05).

**Figure 5.**
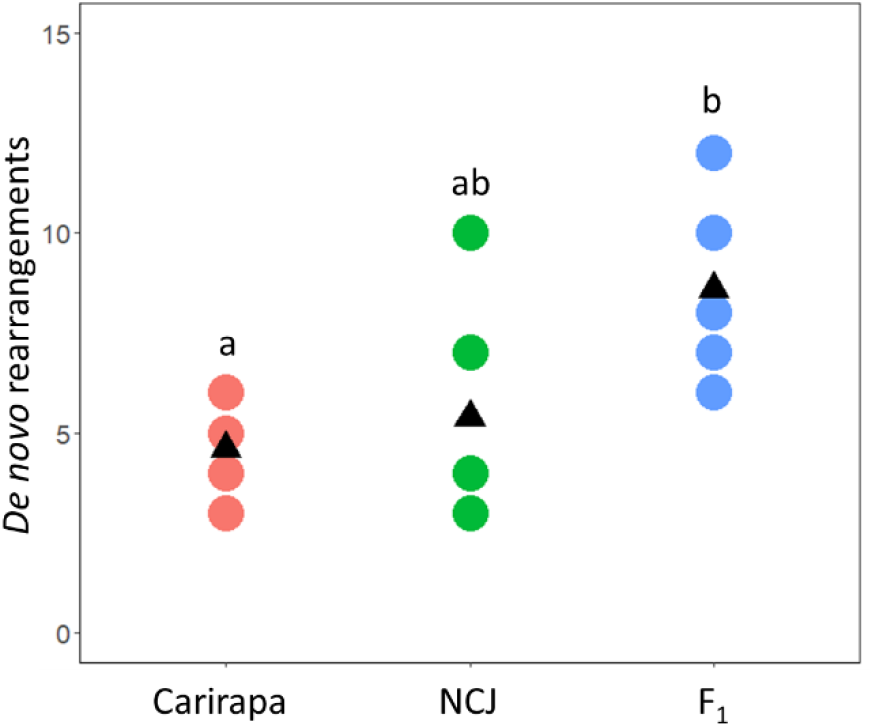
*De novo* rearrangements produced by meiosis in *Brassica* hexaploids carirapa, NCJ and their F_1_ hybrids. Per group, there are 5 different meiosis events. The average per group is drawn with a black triangle. Significance difference are described with letters (TukeyHSD, p< 0.05).

## Discussion

From our study, it is clear that the new F_1_ hybrids produced by the cross between carirapa and NCJ hexaploid lineages were able to tolerate inheritance of different chromosome rearrangements, despite the putative impact of these on meiosis. Initially, we hypothesized that meiosis might be more stable in the F_1_ compared to the parents, as heterozygosity has been linked to crossover frequency in some species (Rowan et al., 2019; Valenzuela et al., 2012), but we observed the opposite. We observed an increase of rearrangements in the F_1_ hybrid, which was significant compared to the carirapa parent (Figure 5). An overall comparison may suggest that the more heterozygous the material, the higher the number of rearrangements that were produced, although further studies are needed to confirm this and to eliminate the confounding effect of the different genotypes. We also found more potential new rearrangements in the B genome of the F_1_ hybrids, mostly affecting alleles inherited from the NCJ parent.

From our analysis, it was also clear that *Brassica* NCJ and carirapa allohexaploids can also accumulate and tolerate multiple structural rearrangements over generations. This occurred even under selective pressure for improved fertility (number of total seeds produced), which hypothetically would be expected to select against aneuploid chromosome complements. This is in contrast to observations of *Brassica napus* synthetic lines in later generations (S_1:11_), where fertility is majorly reduced in aneuploid lines compared to euploid lines (Xiong et al., 2011). Chromosome loss similarly affected fertility in a *Brassica* hexaploid mapping population from carirapa origin, where 30 out of 51 plants were infertile if missing one chromosome (Yang et al., 2018). However, Mason et al. (2014) also found no major effects of chromosome rearrangement on fertility in an NCJ allohexaploid-derived F_2_ population.

The expected chromosome number from the hybrids analyzed in the present study was AABBCC = 2n = 54, but this was rarely observed (Figure 2). In the case of the carirapa lines, two lines had 54 chromosomes but were not euploid. The only NCJ line with a euploid 2n = 54 chromosome complement failed to transmit a copy of chromosome A03 to the resulting F_1_ hybrid, indicating that meiosis was still unstable in this line. Our results may suggest that meiosis in these hybrids is still highly unstable, but that recurrent selection for fertile, viable plants is also acting to eliminate many of these rearrangements, so that plants move from euploid to aneuploid and back again, accumulating minor chromosome rearrangements along the way. Aneuploidy and non-homologous recombination in polyploids and particularly synthetic polyploids are not rare events, and may in some cases even be beneficial due to the potential for creating novel phenotypic variation (reviewed in (Schiessl et al., 2019)).

Non-homologous recombination events reflected known chromosome relationships between the A, B and C genomes, as expected. Most translocations were observed between homoeologous chromosome pairs, as was previously observed in both natural and synthetic *B. napus* (Chalhoub et al., 2014; Samans et al., 2018) and in interspecific *Brassica* hybrids of various types as well as earlier generation NCJ allohexaploid populations (Gaebelein et al., 2019; Mason et al., 2014). For instance, greater changes were observed for homoeologous chromosomes A01/C01 and A02/C02, in accordance with results found in other synthetic *B. napus* lines (Gaeta et al., 2007; Xiong et al., 2011) and natural *B. napus* (Higgins et al., 2018). The B genome was the least affected by fixed genomic rearrangements (Figure 3) and all events observed in this genome were present only in the NCJ allohexaploid types. A possible explanation of why the B genome was more affected in the NCJ lines compared to the carirapa lines is the different method used to generate these hybrid types. The NCJ lines were produced using a two-step crossing method, where initially *B. napus* was crossed to *B. carinata* to form hybrids with genome compositions of 2n = CCAB = 36 (Mason et al. 2012). In these hybrids, the C genomes are present as homologous chromosomes, but the A and B genomes are haploid, and occasionally pair non-homologously with each other or with the C genome (Mason et al. 2010). As well, the preferential loss of the B genome over the A and C genome has been observed in other allohexaploid hybrids (Zhou et al., 2016). In the F_1_ hybrids we observed very few rearrangements involving the B genome. In the F_1_ hybrid parent of population 04, there were two events involving the introgression of B fragments into the C genome, with a carirapa origin. These events were already present as segregating events in the carirapa parent. Based on homoeology between the *Brassica* genomes (Perumal et al., 2020) we identified the most likely position for these introgressions in the C genome. These events were very rare in all of the hexaploids analyzed, but may offer potential for introgression of chromosome fragments from the B genome carrying useful agronomic traits into the A or C genome for *Brassica napus* (rapeseed) crop improvement.

In the different F_1_ hybrids analyzed we observed *de novo* rearrangements involving different chromosomes depending on the population. In some cases, these *de novo* rearrangements occurred close to regions where rearrangements were observed in the parental lines. As an example, in population 08, we observed an increase in the size of the deletion at the top of chromosome C03. This deletion was already present in the carirapa parent, but had a size of ∼6.6 Mb, compared to 16.3 Mb in the F_1_ hybrid (Supplementary Figure 4). This suggests an increase in size for some of the rearrangements, although it was not the rule, as other translocations were directly inherited from the parent without size change. However, already-present translocations may lead to additional irregular pairing during meiosis, as recombinant chromosomes enforce close proximity between homoeologous pairs in translocation heterozygotes, facilitating the production of additional crossover events (Udall et al., 2005) or it might be that those regions in general are more prone to recombine than others (De Muyt et al., 2009). Mwathi et al. (2019) also observed this effect of pre-existing translocations in doubled-haploid NCJ allohexaploid lines, but due to the extreme instability of this population *de novo* translocation events were found to be much more common (2.2 events per plant on average) than events triggered by pre-existing rearrangements (0.8 events per plant on average). In our case, we observed on average 10 *de novo* rearrangements in the F_1_ hybrids, and out of these, 2.8 events on average were proximal to or co-located with a pre-existing event. The difference observed between the two experiments might be attributed to the fact that in our case, the analyzed plants underwent more rounds of meiosis (H3-H7) compared to just two rounds for the Mwathi lines (H_2_) (Mwathi et al., 2019).

Although no fixed pattern of selection was observed across all the allohexaploid lines, several events notably seemed to occur in multiple independent lineages or to be preferentially selected for. This was observed in 3 plants belonging to the N1C2.J1 lineage where the beginning of chromosome A01 was doubled and translocated into C01. The region translocated between A01/C01 has a length of ∼ 2 Mb, and contains the meiosis gene *Cell Division Cycle 20* (*CDC20*). The gene *CDC20* has been found to be crucial in *Arabidopsis thaliana* meiosis, although it might have a different function in *Brassica* (Niu et al., 2015).

The end of chromosome A04 was deleted (∼ 1.1 – 2.5 Mb) in seven plants belonging to three independent lineages (two carirapa and one NCJ). In the case of the NCJ lines, the deletion of this region was already present in the parental *B. napus* cv. “Surpass400_024DH” and in the progenitor line “Surpass” (Higgins et al., 2018). No doubling in the corresponding homoeologous region in C04 was observed for any of these lineages. Although no known meiosis genes fall in this region, the meiosis gene *ASYNAPTIC4* (*ASY4*) is located 2 Mb upstream of the deleted region of A04. In *Arabidopsis thaliana*, this protein is necessary to complete synapsis, cross-over formation and normal localization of interacting proteins ASY1 and ASY3 (Chambon et al., 2018). Interestingly, the homozygous presence of a *ASY3* allele with a tandem duplication region has been associated with more stable autotetraploid *Arabidopsis lyrata* lines, as it causes a reduction in multivalent formation (Seear et al., 2020). The homoeologous region in C04 has recently been associated with fertility in synthetic *B. napus,* where a deletion of the last 1.5 Mb of chromosome C04 reduced seed number (Ferreira de Carvalho et al., 2021). However, this deletion was not associated with a reduction in the number of non-homologous recombination events (Ferreira de Carvalho et al., 2021), suggesting that this effect may only be related to selection for fertility and not to increased stabilization of the genome. The top of chromosome C03 (∼1,2 - 2.3 Mb) was also missing in four plants from the same carirapa lineage (C13), most likely as the result of an event fixed very early on in this lineage, although the four plants differed slightly in the size of the deletion. This A03-C03 region has previously been identified in hexaploid lines as a translocation QTL associated with total seed number in NCJ hexaploids, where the top of chromosome C03 was replaced by a copy of A03 (Gaebelein et al., 2019). Within this region, we find a potential meiotic candidate gene called *BRCA2.* In *Arabidopsis*, the protein BRCA2 interacts in vitro with DMC1 (Dray et al., 2006) and plays a crucial role in homologous recombination (Seeliger et al., 2012).

Five populations from different crosses were analyzed for bias in allele segregation and chromosome inheritance. The B genome was least affected overall, although it was preferentially lost if present as single chromosomes (B05 and B07 in populations 04, 05, and 06). A similar case was observed for the single C09 chromosome (population 05), that was also present less often than expected in one test-cross population. However, this was not an effect seen for all chromosomes present in a single copy: the majority (A03, A04, A05, C01, C02, C06, C08 and C09 in the different populations) were inherited as expected in the progeny (50%). Interestingly, in population 06, there was segregation distortion towards retention of chromosome A09 from the carirapa allohexaploid parent, although both chromosomes were present in the hybrid. This chromosome inherited from the carirapa parent had translocations between chromosomes A09/C09 and A09/C08. Additionally, the chromosome A09 from carirapa had a translocated C09 region of 1.7 Mb, containing the meiosis gene *X-ray induced 1* (*XRI1*). This gene has been shown to be involved in meiosis and post-meiotic stages of male gametes development in plants (Dean et al., 2009). In this F_1_ hybrid, we also just had one C08 chromosome (NCJ), and potentially the retention of an extra A09 occurred as a trisomy compensation of the missing homoeologous region. Compensatory trisomy of chromosomes has been previously described in resynthesized *B. napus* (e.g. trisomy of C05 chromosome, single copy A05)(Xiong et al., 2011). The gain of extra chromosomes to compensate the loss of a homoeologous chromosome can help to prevent further chromosomal instability by providing a partner for the single chromosome (e.g. monosomic-trisomic substitutions) and by preventing the deleterious effects of aneuploidy and changes in gene balance (Xiong et al., 2011).

In populations 05 and 06, the end of chromosome A08 had an allele segregation distortion towards the allele inherited from the carirapa parent. This region was ∼1.3 Mb in size and contained the meiosis genes *ADA2b* and *RAD51D.* Both populations shared the same carirapa and NCJ genotype combination (C13 and N6C2.J2). In rice, *RAD51D* has been shown to be associated with the prevention of non-homoeologous recombination during meiosis (Zhang et al., 2020). The preferential inheritance of an allele putatively related to increased meiotic stability from *B. rapa* over an allele from *B. napus* is unexpected, as *B. napus* is an established allopolyploid species with good meiotic control. However, at least one *B. rapa* line has previously been identified to contain allelic variation relevant for meiotic stability in allohexaploids with *B. carinata* (Gupta et al. 2016). Further confirmation is needed to support this observation.

In population 04, allele inheritance distortion was observed at the end of A07, where alleles from the carirapa parent were preferentially inherited over alleles from the NCJ allohexaploids. The region has a size of ∼5.6 Mb and has also been previously associated with total seed number in an NCJ population (Gaebelein et al., 2019), although it is hard to point to a meiosis candidate gene as several can be found within this interval.

## Author contributions

DQM carried out the experimental analyses, prepared the results figures and tables and drafted the manuscript. ASM and JZ conceptualized the experiment. JZ and JM generated the experimental material. JZ managed all plants in the field and glasshouse, including selection, crossing and self-pollination and DNA extraction. WSZ managed the crossing, self-pollination, harvest in the field and DNA extraction. ASM supervised DQM, and ASM, JB and JZ revised the manuscript. JB managed the generation of SNP array data for the experimental population.

## Acknowledgements

This research was supported by Sino-German DFG Centre grant MA6473/3-1. The authors gratefully acknowledge the technical assistance of Liane Renno.

## Supplementary Figures and Tables

**Supplementary Figure 1: A**. Molecular karyotype and **B**. allele segregation for F_1_ hybrid population 04

**Supplementary Figure 2: A.** Molecular karyotype and **B**. allele segregation for F_1_ hybrid population 05

**Supplementary Figure 3: A.** Molecular karyotype and **B.** allele segregation for F_1_ hybrid population 06.

**Supplementary Figure 4: A.** Molecular karyotype and **B.** allele segregation for F_1_ hybrid population 08

**Dataset S1:** Plant material used in the production of F1 hybrids between carirapa and NCJ hexaploids

**Dataset S2:** Illumina Infinium *Brassica* 90K genotyping array data for the *Brassica* A-, B-, and C- genomes on the *Brassica napus* Darmor-bzh v8.1 and *B. nigra* Ni100 SR reference genomes for *Brassica* control species, *Brassica* hexaploid “NCJ”, “Carirapa”, F1 hybrids, test-cross parents, and test- cross populations. Allele AA, BB = homozygous, AB = heterozygous, and NC = not-called.

**Dataset S3**: Illumina Infinium *Brassica* 90K SNP array logR ratio data for the *Brassica* A-, B-, and C- genomes on the *Brassica napus* Darmor-*bzh* v8.1 and *B. nigra* Ni100 SR reference genomes for *Brassica* control species, *Brassica* hexaploid “NCJ”, “Carirapa”, F1 hybrids, test-cross parents, and test- cross populations.

